# Enhanced envelope coding following acoustic trauma is detrimental to neural coding of speech in noise

**DOI:** 10.1101/2022.03.16.484675

**Authors:** Satyabrata Parida, Michael G. Heinz

## Abstract

People with hearing loss struggle understanding speech in noisy backgrounds. Speech-intelligibility models highlight the importance of slow-varying envelopes of speech and noise; however, the physiological basis of impaired perception remains unclear. We provide neurophysiological evidence that, although acoustic trauma enhances both speech and noise envelopes, disruptions in cochlear tonotopicity preferentially enhance neural responses to noise in noisy speech. These results provide mechanistic insights into everyday-communication challenges created by hearing loss.

---

Listeners with noise-induced hearing loss (NIHL) struggle to understand speech in noisy environments despite having little difficulty in quiet environments. These deficits most severely manifest in fluctuating-noise environments, such as in restaurants amidst other talkers, and are not resolved even using state-of-the-art hearing aids^1^. Hearing aids, which are fit based on the audiogram (clinical gold-standard), provide limited benefit in these noisy environments because audiograms measure hearing sensitivity in quiet conditions and do not reflect suprathreshold deficits in daily communication. A better understanding of the neural coding of real-world sounds following hearing loss will guide strategies to mitigate suprathreshold hearing difficulties, which will ultimately lead to better clinical outcomes.

One such suprathreshold feature is the slowly varying envelope of speech, which is critical for speech perception^2–4^. Previous neurophysiological studies have shown an enhancement in envelope coding of various narrowband and broadband signals, including speech and noise when presented alone following NIHL^5–8^. However, the effects of NIHL on speech and noise envelopes when presented simultaneously (as in real-world scenarios) are unknown. Psychoacoustic studies suggest a detrimental role of enhanced envelope coding on speech-in-noise representation^9^, but this hypothesis has not been tested in impaired neural responses.

We recorded spike-train data from single auditory-nerve (AN) fibers of anesthetized male chinchillas that either had normal hearing (NH) or mild-to-moderate hearing loss (Fig. S1), the most clinically prevalent degree of hearing loss^10^. As expected, AN fibers from NIHL animals had elevated threshold, broader bandwidth, and reduced tip-to-tail ratio (TTR) in frequency tuning curves, as well as reduced spontaneous rate^11–13^. Spike-train data were recorded in response to a naturally uttered speech sentence (*S*), a stationary speech-shaped noise (SNN; Fig. 1B), and a fluctuating noise that had 8-Hz sinusoidal amplitude modulation (FLN; Fig. 1C). Spike-train data were also collected in response to noisy-speech mixtures (*SN*) at three (−10, -5, and 0 dB) different perceptually relevant signal-to-noise ratios (SNRs) that were chosen to leverage the substantial difference in speech intelligibility across maskers and between listeners with and without hearing loss^14^. Specifically, NH listeners (but not listeners with hearing loss) can take advantage of fluctuations in maskers within this SNR range. Envelope coding was significantly enhanced for HI AN-fiber responses to all stimulus types (Fig. S2), extending previous reports of post-NIHL envelope enhancement to several new stimuli (natural speech, SNN, and FLN). Notably, this enhancement was greater for FLN than for SNN.

**Figure 1.**
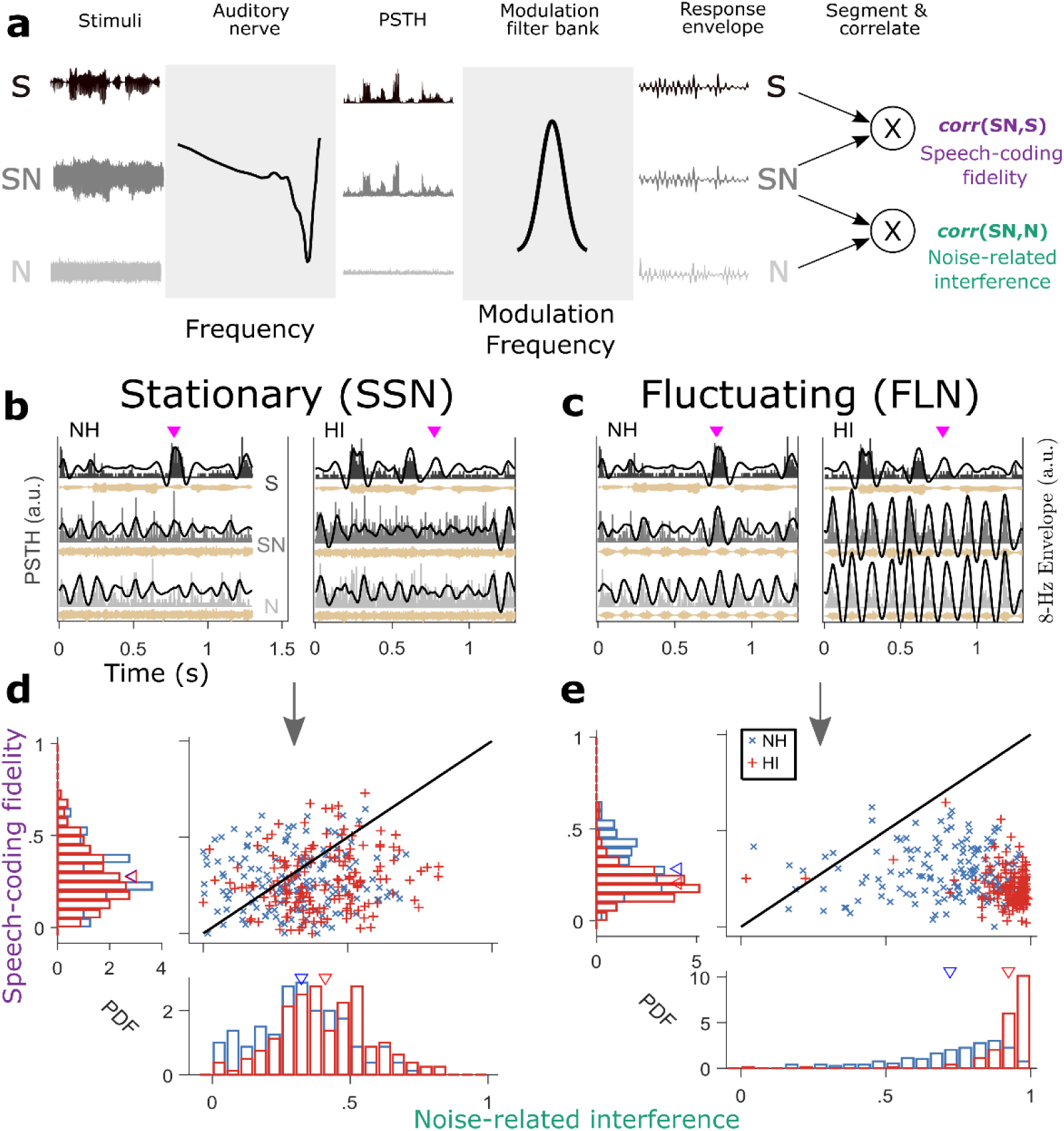
Exemplar neural data highlight the deleterious effects of a fluctuating masker on speech-in-noise envelope coding for the hearing-impaired group, consistent with perception. **a**, Framework to quantify *speech-coding fidelity* [*corr*(*SN, S*)] and *noise-related interference* [*corr*(*SN, S*)]. **b-c**, Exemplar PSTHs from one NH (CF = 4.7 kHz) and one HI AN fiber (CF = 4.6 kHz) for SNN (**b**) and FLN (**c**) at -5 dB SNR. Black trace = 8–Hz envelope. While the NH fiber responded strongest to a (high-frequency) fricative (magenta triangles near 0.8 s) in *S* (as expected based on tonotopic coding), the HI fiber over-responded to (low-frequency) voiced speech (0.3 – 0.7 s) and responded weakly to the fricative. Furthermore, HI-fiber envelopes are enhanced relative to the NH fiber, particularly for the fluctuating-masker condition (both SN and N). **d-e**, Probability density functions (PDF) for *corr*(*SN, S*) and *corr*(*SN, S*) for SNN (**d**) and FLN (**e**) at 0-dB SNR for the 8-Hz modulation filter. Triangles represent group means. Noise-related interference was substantially higher for the HI group, especially for the fluctuating masker. NH, normal hearing; HI, hearing impaired; a.u., arbitrary units; CF, characteristic frequency; PSTH, peristimulus time histogram.

To test whether these individual enhancements were detrimental to overall speech-in-noise coding, the relative contributions of speech (*S*) and noise (*N*) to noisy speech (*SN*) responses were quantified using a speech-intelligibility model^15^ (Fig. 1). In this framework, AN spike trains were used to construct peristimulus time histograms (PSTHs), which were further filtered by a modulation filter-bank that mimics properties of midbrain neurons^16^. The center frequencies of these modulation filters spanned 4 to 32 Hz, frequencies most important for speech perception^17^. Speech contributions to *SN* responses, termed *speech-coding fidelity*, were quantified as the correlation between response envelopes of *SN* and *S* [*corr*(*SN, S*)]. Similarly, noise contributions to *SN* responses, termed *noise-related interference*, were quantified as the correlation between *SN* and *S* response envelopes [*corr*(*SN, S*)]. Note that *corr*(*SN, S*) quantifies the salience of unwanted distracting modulations due to background noise (known as modulation detection interference^18^), and captures important aspects of speech-in-noise coding that are complementary to *speech-coding fidelity* captured by *corr*(*SN, S*). Speech-coding fidelity and noise-related interference were estimated for individual AN fibers at each modulation frequency. These estimates were used to construct probability density functions (PDFs) for each hearing-status group (e.g., Fig. 1d-e).

For each correlation metric, a diagnostic d-prime was used to quantify the distance between the HI and NH group PDFs (Fig. 2a). d-prime for *corr*(*SN, S*) was often negative (especially for negative SNRs), indicating poorer speech-coding fidelity for the HI group compared to NH, especially for FLN. Surprisingly, d-prime for *corr*(*SN, S*) was always positive, indicating more severe noise interference for the HI group, especially for FLN.

**Figure 2.**
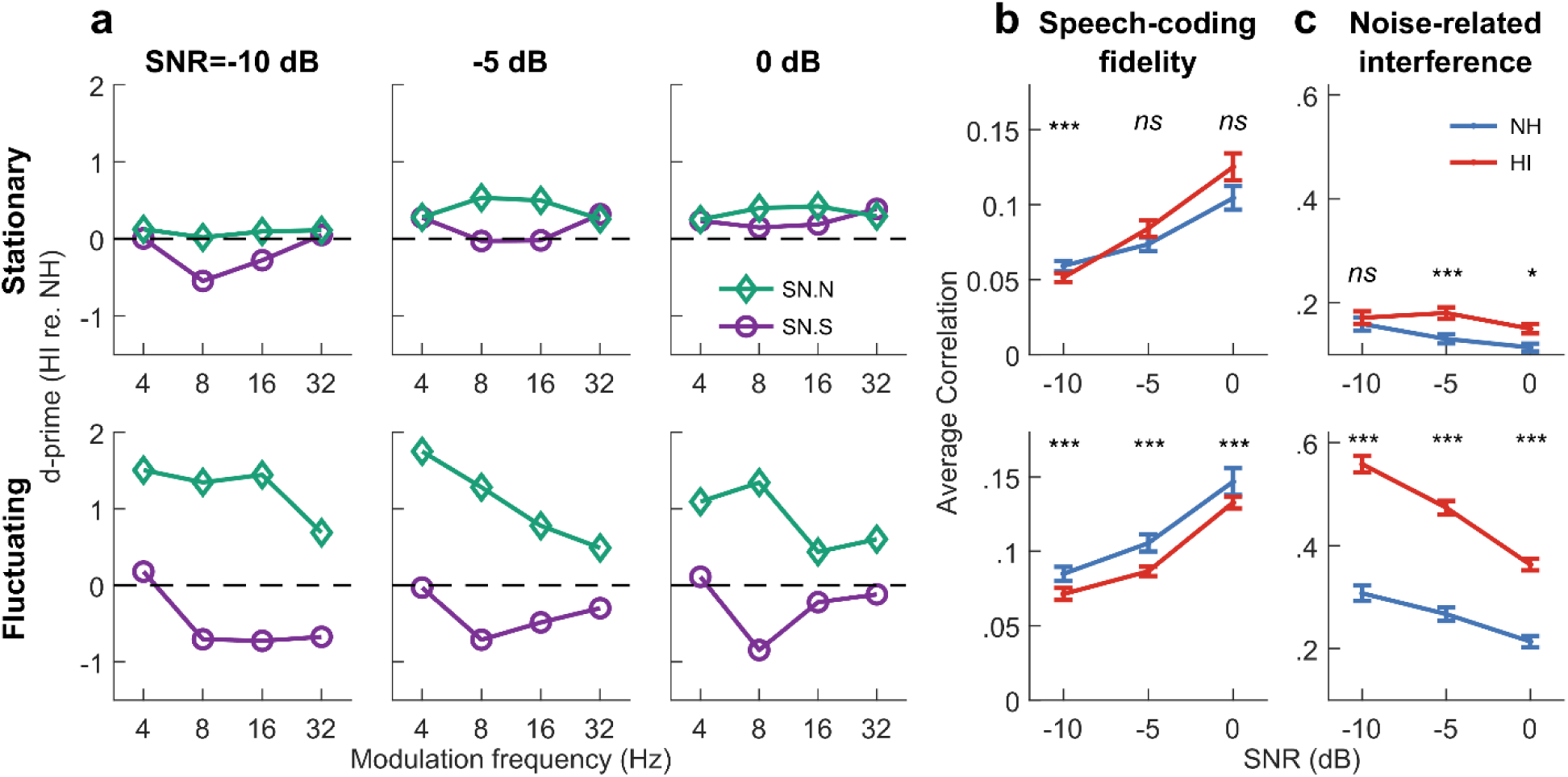
Noise-related envelope was preferentially enhanced following acoustic trauma in neural responses to noisy speech, especially for the fluctuating masker. **a**, A diagnostic d-prime metric indicates that *corr*(*SN, S*) was regularly enhanced (>0) following NIHL, whereas *corr*(*SN, S*) was sometimes degraded (<0). **b**, Speech-coding fidelity was reduced for the HI group, particularly for the fluctuating masker. **c**, Noise-related interference was significantly enhanced for the HI group, especially for the fluctuating masker. This enhanced *corr*(*SN, S*) highlights the distracting nature of fluctuating noise, the condition where listeners with NIHL struggle the most during everyday communication. Error bars represent s.e.m.

Another way of considering the perceptual relevance of these data is to estimate the average (*rms*) correlation across modulation frequencies for each group, which can be treated as a neural estimate of speech intelligibility^15^ (Fig. 2b-c). These results reiterate the same key points as in Fig. 2a, i.e., that: (1) speech-coding fidelity was poorer and noise-related interference was greater for the HI group, and (2) these degradations were worse for FLN than SNN. These neural results are strikingly parallel to psychoacoustic results where listeners with hearing loss struggle more in fluctuating-masker conditions than in stationary-masker conditions^14^.

Although consistent with psychoacoustics, these results are counterintuitive as speech and noise are simply two signals to the cochlea, and an asymmetric enhancement of one signal in response to their mixture is unexpected. What mechanism is responsible for this asymmetric enhancement of *N*? The usual suspect is broadened frequency-tuning bandwidth, which regularly accompanies NIHL^12,19^ (Fig. S1d). Alternatively, our recent studies show that disruption in cochlear tonotopy, the functional connectivity between sound frequency and cochlear place, which occurs following NIHL can play a key role in speech-in-noise coding^20–22^. To evaluate the relative contributions of these two factors (i.e., bandwidth and tonotopy), a toy model was created. The stimuli input to the model were a swept narrowband “signal” (*Ŝ*) and a broadband noise 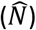, which mimic key spectrotemporal properties of speech (spectrally sparse over short time scales) and noise (broadband), respectively (Fig. 3a-b). Model responses to 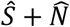 (at -5 dB SNR) were also simulated. Responses of three model AN fibers were investigated: a normal-hearing fiber (narrow bandwidth and high TTR), an impaired fiber with broad bandwidth but high TTR, and an impaired fiber with narrow bandwidth but low TTR (Fig. 3c-d). For these systems, 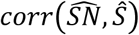 and 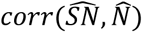 were estimated to quantify signal-coding fidelity and noise-related interference. Results showed that only the low-TTR system displayed degradations similar to those observed in our neural speech-in-noise data [e.g., >50% increase in *corr*(*SN, S*) following HL, Fig. 2c]. Overall, temporal responses of the low-TTR model fiber replicated the key results in the neural data: (1) a reduced-TTR system over-represents low-frequency energy, thus rendering responses to be non-tonotopic, (2) 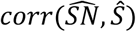 is weakly affected suggesting that 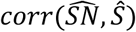 is driven by common non-tonotopic (low-frequency) components in 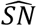 and *Ŝ*, and (3) 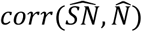 is substantially enhanced, suggesting that noise with substantial low-frequency energy can be particularly distracting.

**Figure 3.**
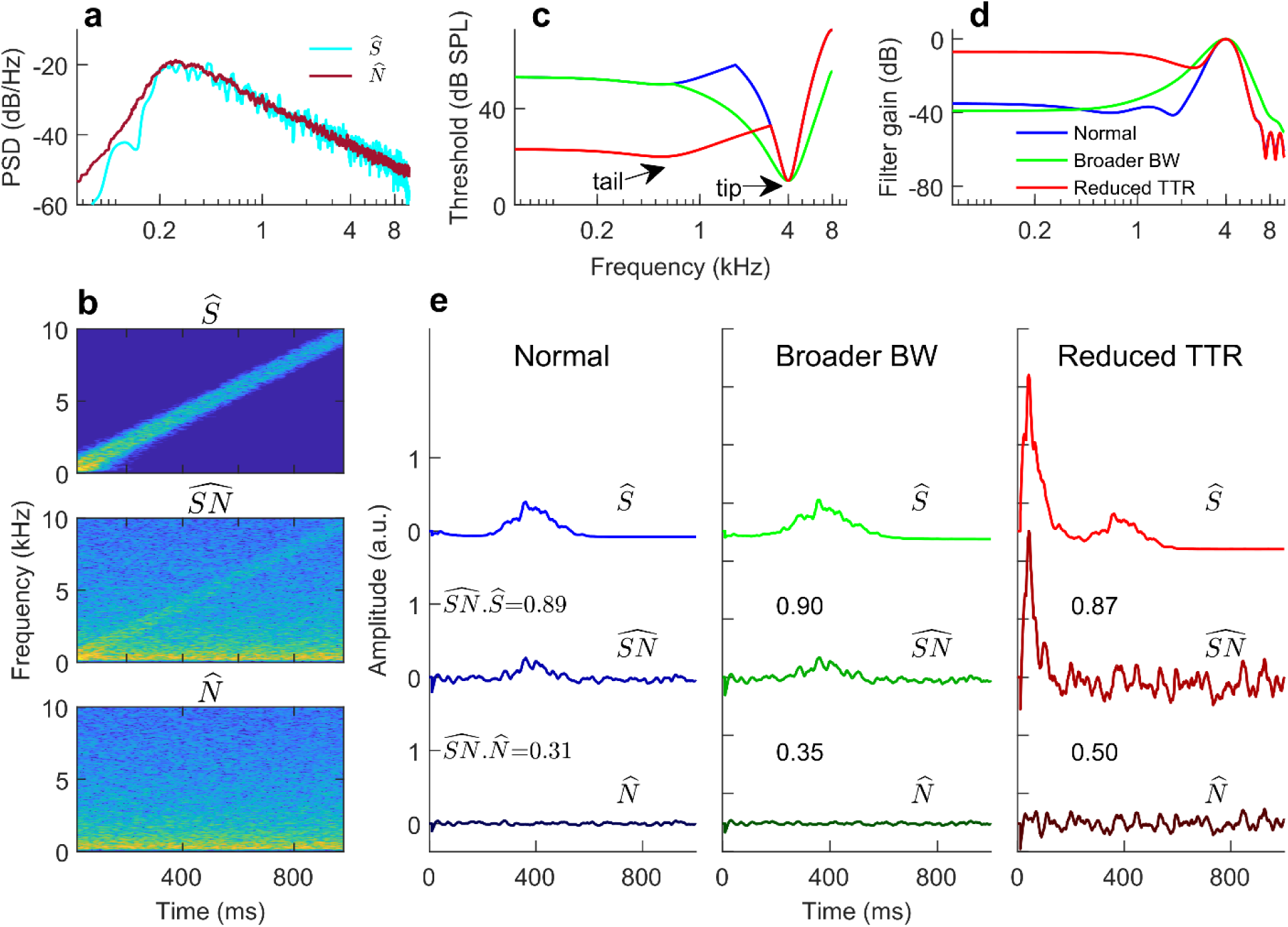
Toy model suggests that the preferential enhancement of noise-related envelopes results from distorted tonotopy following NIHL. **a–b**, The two signals used in the toy model had similar long-term spectrum (**a**), but different short-term spectrotemporal properties (**b**). Over short time scales, the swept narrowband “signal” (*Ŝ*) is bandlimited but 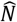 is broadband. 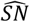 was generated by mixing *Ŝ* and 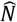 at -5 dB SNR. **c**, Frequency-tuning curves for three model AN fibers that help disentangle the effects of broader tip bandwidth from distorted tonotopy (i.e., reduced TTR). **d**, Transfer functions for the model fibers. **e**, Response envelopes of the model fibers (with correlation values shown as in-panel text) suggest that distorted tonotopy is more detrimental to speech-in-noise coding than broader bandwidth (BW).

In summary, these results show that enhanced envelope coding following acoustic trauma is detrimental to the neural coding of speech-in-noise. This degradation was particularly strong in fluctuating backgrounds, consistent with the inability of listeners with NIHL to benefit from masker fluctuations^14^. These results also emphasize the importance of considering both tonotopic speech-coding fidelity and enhanced noise-related interference in accounting for speech perception by listeners with hearing loss (e.g., in speech-intelligibility models)^23^. Because distorted tonotopy, the degree of which varies across hearing-loss etiologies^21^, is the dominant contributor to these degradations in speech-in-noise coding following NIHL, this understudied suprathreshold deficit likely contributes to individual differences in speech perception among listeners with sensorineural hearing loss. Further studies are warranted to establish the relation between speech perception and noninvasive assays of distorted tonotopy^22^, especially in real-world listening conditions since environmental noises have substantial low-frequency energy, which can exacerbate the effects of distorted tonotopy^20,22^.

## Methods

All experimental procedures followed PHS-issued guidelines and were approved by Purdue Animal Care and Use Committee (Protocol No: 1111000123). Data were collected from young (< 1 year old, weighing between 400 and 700 g) male chinchillas (189 AN fibers from 12 normal-hearing chinchillas; 132 AN fibers from 7 hearing-impaired chinchillas) using standard procedures in our laboratory^5,6,21^. Animals were socially housed in pairs until they went through a procedure involving anesthesia. Housing was maintained in a 12–hour light /12–hour dark cycle. All animals received daily nonauditory environmental enrichment (chewing toys and dietary treats).

### Noise exposure and electrophysiological recordings

Animals were exposed to 116 dB SPL (C-weighted) octave-band noise centered at 500 Hz for two hours using an enclosed subwoofer (Selenium 10PW3, Harman), which was placed ∼30 cm above the animal’s head. Exposure level was calibrated near the animal’s ear canal using a sound-level meter (886–2, Simpson, Elgin, IL, USA). Animals were anesthetized using xylazine (2 to 3 mg/kg, subcutaneous) and ketamine (30 to 40 mg/kg, subcutaneous) prior to their noise exposure. Animals’ vital signs were monitored throughout all procedures using a pulse oximeter (Nonin 8600V, Plymouth, MN). An oxygen tube was placed near the animals’ nostrils. A xylazine-reversal agent (Atipamezole, 0.4 to 0.5 mg/kg, intraperitoneal) was used after the procedure to facilitate speedy recovery from anesthesia. Animals were given warm lactated Ringer’s solution (before and after the procedure, 6 cc each, subcutaneous) and a high-calorie diet (after the procedure for three days, DietGel Criticare, ClearH20, Portland, ME, US). Animals were allowed at least a two-week recovery period following noise exposure before any electrophysiological recordings were obtained. The animal’s rectal temperature was maintained at 37 °C using a feedback-controlled heating pad (50-7053F, Harvard Apparatus) and the room temperature was kept elevated at 24°C during any anesthetic procedure.

Animals were screened with hearing assessments before and after (the two-week recovery period) noise exposure. The same anesthetic and recovery procedures were followed as noise exposure. A transducer-microphone pair (Etymotic ER-2, Etymotic ER-10B, Etymotic Research, Elk Grove Village, IL, USA) was used for acoustic calibration and sound presentation with a foam ear tip inserted into the external ear canal.

Auditory brainstem responses (ABRs) were recorded using subdermal needle electrodes in a vertical montage (differential mode; active electrodes near the vertex and mastoid and common ground near the nose). ABRs were band-pass filtered (0.3–3 kHz, ×20,000 gain) using analog filters/amplifiers (ISO-80, World Precision Instruments, Sarasota, FL; SR560, Stanford Research Systems, Sunnyvale, CA; 2400A, Dagan, Minneapolis, MN). ABRs were collected in response to tone pips (5-ms duration, 0.5-ms on and off ramp, 31-ms repetition period, 500 repetitions per polarity) at five frequencies (0.5, 1, 2, 4, and 8 kHz) from 0 to 80 dB SPL in 10 dB steps. ABRs for another intensity (odd multiple of five) near the preliminary threshold estimate were also collected to fine-tune final threshold estimate. Threshold was estimated based on a cross-correlation analysis^22,24^. Briefly, the ABR at a high intensity (60/80 dB SPL for normal-hearing/noise-exposed animals) was used as the template and was cross-correlated with ABRs at lower intensities (response correlation) or physiological noise in the same session (correlation noise floor). Threshold was estimated as the intensity at which linear regression of response correlation crossed three standard deviations above the correlation noise floor.

Distortion product otoacoustic emissions (DPOAEs) were collected in response to tone-pairs at *f*_1_ (at 75 dB SPL) and *f*_2_ (at 65 dB SPL, ranging from 0.5 to 12 kHz) with *f*_2_/*f*_1_ = 1.2 using the same transducer-microphone pair. DPOAE level was defined as the peak (in dB SPL) at 2*f*_1_ − *f*_2_ in the response spectrum.

### Surgical preparation

Anesthesia was induced using the same xylazine/ketamine doses as for noise exposure. A tracheostomy was performed to allow low-resistance air pathway to minimize breathing-related acoustic artifacts. The cartilaginous portions and muscles near the external ear were dissected to allow insertion of custom-made hollow brass ear bars into the external auditory meatus. After stereotaxic positioning of the animal, a posterior fossa approach was used for craniotomy until the brainstem was visible, following which, cotton pellets were used to push the brainstem away from the lateral wall to reveal the exit of the 8^th^ cranial nerve from the internal auditory meatus. The posterior bulla was vented with a 30-cm long polyethylene tube to maintain middle-ear pressure. High impedance (10–50 MΩ) glass micropipettes filled with 3M NaCl were used as electrodes. Animals were supplemented with lactated Ringer’s throughout the experiment (∼1 ml/hour). The room temperature, additional oxygen, the animal’s rectal temperature and vitals were maintained as described previously. Experiments usually lasted for 18–24 hours and terminated if sudden changes in frequency-tuning-curve thresholds were detected for two or more consecutive auditory-nerve fibers, following which a lethal barbiturate dose (2 cc intraperitoneal, Euthasol, Virbac Corporation, Westlake, TX) was administered.

### Neurophysiological recordings and stimuli

Recordings were amplified (2400A, Dagan Corporation, Minneapolis, MN) and filtered (0.03 to 6 kHz; 3550, Krohn-Hite Corporation, Brockton, MA), and isolated spikes were identified using an amplitude-window discriminator (BAK Electronics, Mount Airy, MD, USA). Single fibers were identified by looking/listening for sound evoked activity to broadband noise as the search stimulus (∼20 dB re 20 μPa/ √Hz for normal-hearing animals, and higher as required for hearing-impaired animals) while advancing the electrode in 2-3 μm steps. When a fiber was encountered, an automated frequency tuning curve (FTC) was generated^25^, followed by routines to estimate its spontaneous rate (over a 30-s silence period) and rate-level function at its CF (CF estimated from the FTC). The CF for individual fibers was estimated as the local minimum closest to the high-frequency-side slope of FTCs; this approach offers a close estimate of the CF for an auditory-nerve fiber appropriate for its cochlear location^26^. Local 10-dB quality factor or *Q*_10_ was estimated as the 10-dB bandwidth relative to the threshold at CF. For W-shaped FTCs, the narrowest 10-dB bandwidth near CF was considered for local *Q*_10_, which is similar to psychoacoustic approaches for estimating tuning^27^ and disentangles broadened bandwidth effects from distorted tonotopy^20^.

Next, SNR-specific routines were employed to collect spike-train data for relevant speech, noise, and noisy-speech files. A naturally spoken sentence was used as the speech stimulus (list #3, sentence #1 of the Danish speech intelligibility test)^28^. Two types of noises were used: steady-state noise (SNN) and 8-Hz sinusoidally amplitude modulated (fluctuating) noise (FLN). Both noises were frozen (i.e., a single instance was used for all AN fibers) and were spectrally matched to ten sentences spoken by the same speaker as the speech sentence. Overall speech level was set to 65- and 80-dB SPL for normal-hearing and hearing-impaired chinchillas, respectively. Noise was scaled and added to speech to achieve a desired SNR (either -10, -5, or 0 dB). The order of SNR conditions during the experiment was pseudo-random. For each SNR-specific routine, stimuli (speech, noises, and their mixtures) were presented in an interleaved manner. Following data collection, spike data were screened to remove any (artifactual) spikes that rarely (< 0.1% for most units) occurred before the absolute refractory period (0.6 ms) following the previous spike.

### Envelope-correlation analyses

Components of speech and noise in responses to noisy speech were quantified using correlational analyses on the response envelopes using a multi-resolution framework^15^. Recorded AN-fiber spike trains were used to construct peristimulus time histograms (PSTH) with 0.5 ms bin resolution. This PSTH was processed through a modulation filter bank (four filters) with center frequencies = 4, 8, 16, and 32 Hz. Filters were fourth order, octave wide, and zero phase. Contributions of speech and noise to noisy-speech responses were quantified using correlational analyses. Envelope correlation between responses to speech (*S*) and noisy speech (*SN*) is termed *speech-coding fidelity* and is denoted by *corr*(*SN, S*). Similarly, envelope correlation between responses to noise (*N*) and noisy speech is termed *noise-related interference* and is denoted by *corr*(*SN, S*). These correlation metrics were estimated as follows.

For each AN fiber (*f*_*c*_) and modulation filter (*f*_*m*_) combination for each noise condition, the output of the modulation filter for all stimuli (i.e., *S, N*, and SN) were divided into segments of 2/*f*_*m*_, and correlation metrics [i.e., *corr*(*SN, S*) and *corr*(*SN, S*)] were estimated for each segment. These segment correlation values were rectified (i.e., set to zero if negative). The final correlation value for that AN fiber at that modulation frequency was estimated as the average of the rectified correlation values across all segments. For each group, these final correlation values were pooled to construct probability density functions at each modulation frequency per noise condition (e.g., Fig. 1d-e). Only AN fibers with *f*_*c*_ < 5 kHz were considered to avoid major sampling biases in *f*_*c*_ between groups.

Group differences were quantified from the distance between these correlation distributions (Fig. 2a) at each modulation frequency, estimated by the following diagnostic d-prime metric:

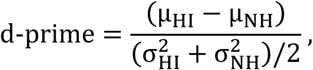

where *μ* and *σ* denote the mean and standard deviation of the probability density function. Note that negative d-prime for *corr*(*SN, S*) indicates poorer *speech-coding fidelity* for the HI group. Similarly, positive d-prime for *corr*(*SN, S*) indicates greater *noise-related interference* for the HI group.

Average correlation values in Fig. 2b-c were estimated as the root-mean-square (*rms*) of the correlation value across *f*_*m*_ for each AN fiber, as done in speech-intelligibility modeling^15,29^. Mean and standard error for each group are based on these final correlation values across AN fibers.

### AN-fiber modelling

To disentangle the effects of broader tuning bandwidth from distorted tonotopy, a toy model was created with simplified stimuli and model AN fibers. Two signals, a broadband noise 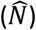 and a swept narrowband-noise signal (*Ŝ*), both with *red long-term spectrum* (i.e., 1/*f*^2^ spectrum) were used as stimuli to mimic the spectrotemporal properties of noise (broadband over short time scales) and speech (narrowband over short time scales), respectively. Red spectrum was used because it is closer to the long-term spectrum of speech thank pink or white noise^30^. *Ŝ* was designed by first generating red noise, then applying frequency demodulation (linear trajectory from 200 Hz to Nyquist frequency – 200 Hz) and low-pass filtering (bandwidth = 400 Hz, fourth order), and finally applying frequency modulation (to invert the initial frequency demodulation)^31^. Thus, *Ŝ* was a swept narrowband noise with a bandwidth of 400 Hz at any single instance. 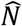 was 5 dB higher in level than *Ŝ*; therefore, signal to noise ratio for 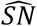 was -5 dB. All signals were high-pass filtered at 250 Hz.

Responses of three model AN fibers were investigated: (1) a normal-hearing fiber with a narrow bandwidth (*Q*_10_ = 2.3) and high (40 dB) TTR, (2) an impaired fiber with a broad bandwidth (*Q*_10_ = 1.5) but high (40 dB) TTR, and (3) an impaired fiber with a narrow bandwidth (*Q*_10_ = 2.3) but low (10 dB) TTR. To design the system function for each model fiber, first an FTC was designed with a tip (fourth order gammatone filter) and a tail (first order gammatone filter centered at 500 Hz with five times the equivalent rectangular bandwidth for humans at 500 Hz^27^). System transfer functions were derived from inverted FTCs with maximum gain set to 0 dB. Filtered signals were half-wave rectified. Response envelope was obtained by lowpass filtering this rectified output (cut-off = 16 Hz, fourth order, zero phase). Response envelope correlation was estimated for this one envelope filter because of the simplified nature of *Ŝ*.

### Statistical analysis

Statistical analyses were performed in R (version 4.0.3) using linear mixed-effects models (lme4 package^32^) to report the effects of *group* (normal-hearing or hearing-impaired). Reported p- and F-values are based on Type II Wald F tests^33^. Log-transformed CF was included in all statistical models for AN data. CF was log-transformed because of the approximate logarithmic spacing of frequency in the cochlea. A p-value of 0.05 was used as the threshold for significance. Significance codes in Fig. 2: 0 ‘***’ 0.001 ‘**’ 0.01 ‘*’ 0.05 ‘*ns*’ 1. In Fig. S2, statistical analysis of envelope coding for noise-alone is based on a single model consisting of group, noise type (SNN or FLN), and SNR as the fixed effects, and AN fiber identifier as a random effect; interactions were included between fixed effects and dropped when not significant (p > 0.05) in order of decreasing p-value.

## Acknowledgement

We thank Vibha Viswanathan, François Deloche, and Ivy Schweinzger for providing valuable feedback on earlier versions of the manuscript. This work was supported by the National Institute of Health (R01-DC009838) and an International Grant (G72) from Action in Hearing loss (UK).

## Author contributions

All authors contributed to the experimental design and writing of the paper. SP performed the experiments and formal analyses. MGH acquired funding.

## Supplemental Information

**Figure S1.**
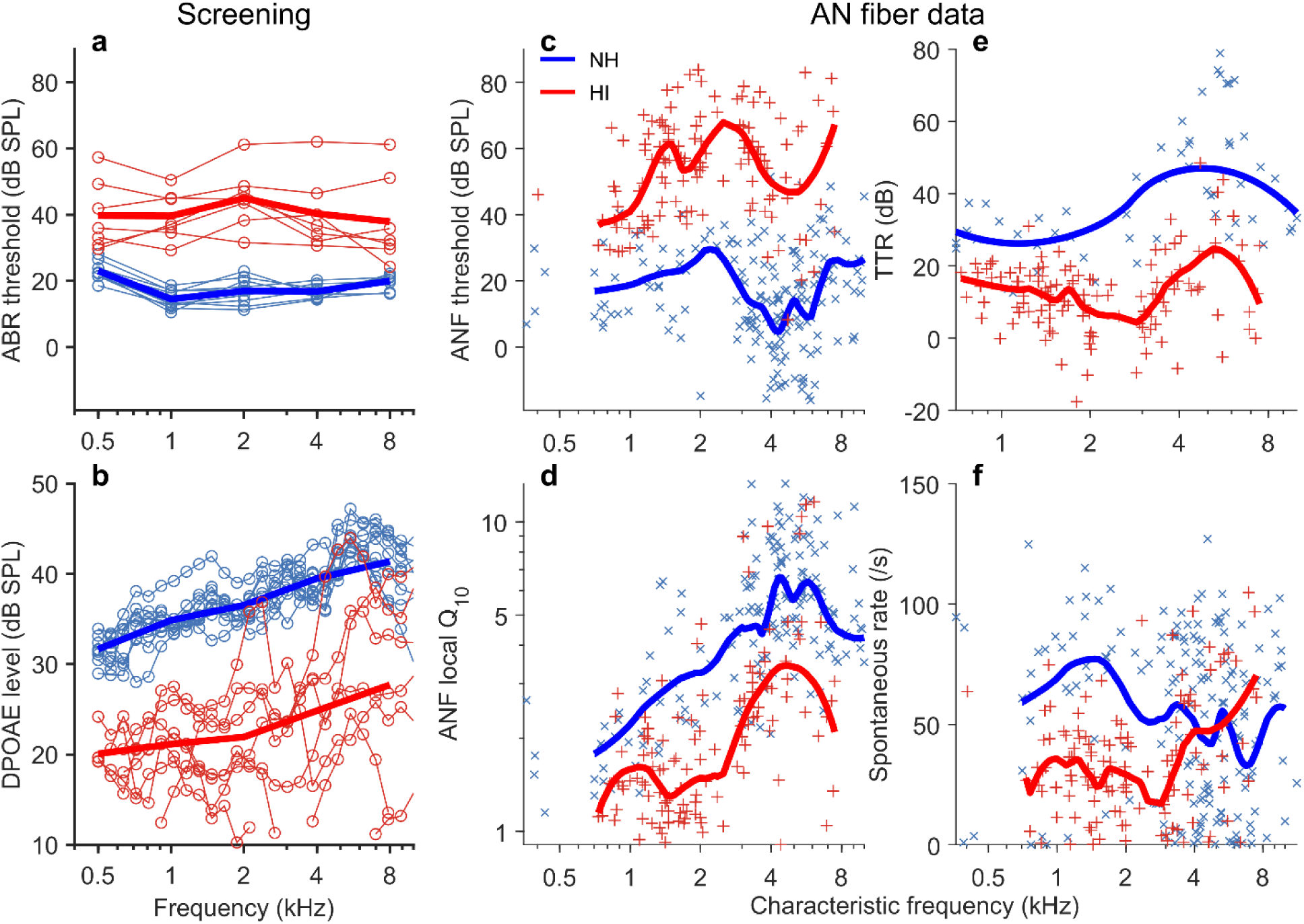
Acoustic trauma caused mild-moderate hearing loss resulting in reduced audibility, broadened frequency tuning, and reduced tonotopicity. **a**, Auditory brainstem response (ABR) thresholds were elevated on average by ∼20 dB for the HI animals (main effect of group, F = 252.8, p < 2.2×10^−16^). Thin lines with symbols represent individual animals (n=9/7, NH/HI); thick lines represent group averages. **b**, Similarly, distortion product otoacoustic emission (DPOAE) levels were decreased, which indicates substantial outer-hair-cell damage (F = 1153.8, p < 2.2×10^−16^). **c-f**, For auditory-nerve fibers, frequency-tuning curve (FTC) threshold was elevated (**c;** F = 68.3, p = 8.5×10^−7^), frequency tuning was broadened near the FTC-tip as quantified by local *Q*_10_ (**d;** F = 11.6, p = 3.0×10^−3^). Tip-to-tail ratio (TTR), which is the difference in threshold at the tail and the tip, was also consistently reduced (**e;** F = 111.4, p = 1.7×10^−9^). Trend lines were computed by locally robust loess regression (smoothing window span = 40%). Spontaneous rate was also reduced for the HI group (**f;** F = 8.7, p < 8.5×10^−3^), suggesting substantial inner-hair-cell dysfunction^13^. ANF, auditory nerve fiber.

**Figure S2.**
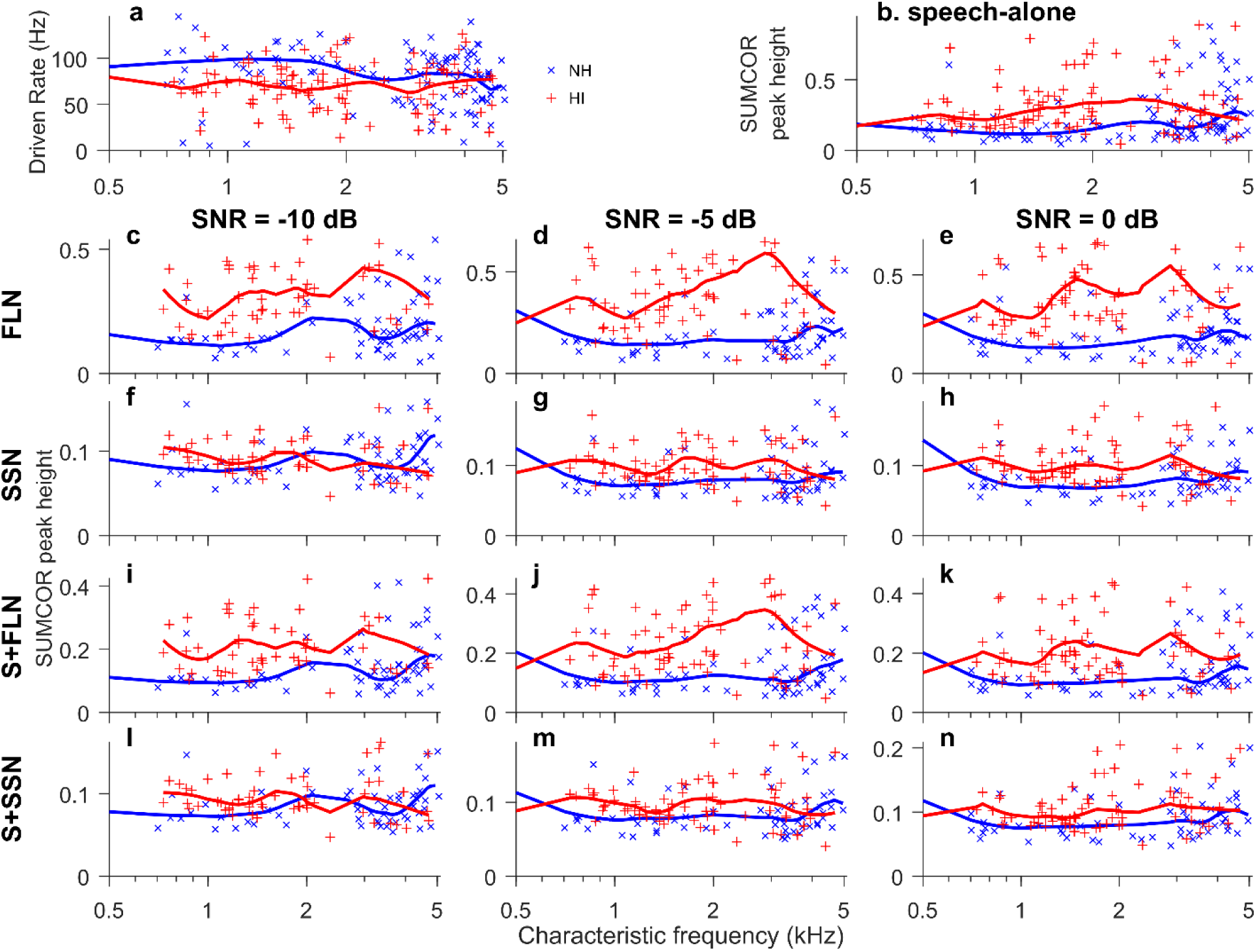
Envelope coding was significantly enhanced for speech, noise, and noisy speech following NIHL, especially for the fluctuating masker. **a**, Driven rate was significantly (group, F = 10.98, p = 0.011) reduced following hearing loss. **b**, Sumcor peak height, a metric for envelope-coding strength, shows significant envelope enhancement for the hearing-impaired (HI) group for speech-alone but only at lower frequencies (CF < 3.5 kHz: group, F = 6.4, p = 0.012; CF < 5 kHz: group, F = 0.92, p = 0.34). **c-h**, Same format as **b**. Envelope coding was also enhanced for noise-alone stimuli, including both FLN and SNN (group, F = 28.1, p = 2.5×10^−7^). Envelope coding was particularly enhanced for the fluctuating noise (group × noise type, F = 224.8, p < 2.2×10^−16^; noise, F = 1191, p < 2.2×10^−16^). **i-n**, Same format as **b**. Envelope coding was also significantly enhanced for noisy-speech stimuli, particularly for S+FLN (group, F = 6.3, p = 0.012; group × noise, F = 102.2, p < 2.2×10^−16^, noise, F = 620.2, p < 2.2×10^−16^). Sumcors were filtered between 4/√2 Hz to 32×√2 Hz to emphasize important modulation bands for speech (same bandwidth as in Fig. 2). Trend lines were computed by locally robust loess regression (smoothing window span = 40%). NIHL, noise-induced hearing loss; S, speech-alone; SNN, stationary noise; FLN: 8-Hz sinusoidally amplitude modulated (fluctuating) noise.

